# CENsible: Interpretable Insights into Small-Molecule Binding with Context Explanation Networks

**DOI:** 10.1101/2023.10.18.562959

**Authors:** Roshni Bhatt, David Ryan Koes, Jacob D. Durrant

**Affiliations:** Department of Computational and Systems Biology, University of Pittsburgh, Pittsburgh, PA 15260; Department of Biological Sciences, University of Pittsburgh, Pittsburgh, PA 15260

**Author notes:** Correspondence: Jacob Durrant, Department of Biological Sciences, University of Pittsburgh, Pittsburgh, PA 15260.

## Abstract

We present a novel and interpretable approach for predicting small-molecule binding affinities using context explanation networks (CENs). Given the specific structure of a protein/ligand complex, our CENsible scoring function uses a deep convolutional neural network to predict the contributions of pre-calculated terms to the overall binding affinity. We show that CENsible can effectively distinguish active vs. inactive compounds for many systems. Its primary benefit over related machine-learning scoring functions, however, is that it retains interpretability, allowing researchers to identify the contribution of each pre-calculated term to the final affinity prediction, with implications for subsequent lead optimization.

## Introduction

Structure-based computer-aided drug discovery (CADD) leverages computer algorithms to design and discover new drug candidates, with the twin goals of accelerating early-stage drug design and reducing costs. Among CADD techniques, molecular docking is particularly popular. First, a docking program predicts how small-molecule ligands (e.g., candidate drugs) position themselves within a protein target’s binding pocket (i.e., the 3D geometry of binding, or “binding pose”). Second, a docking scoring function maps those binding geometries to scores correlating with efficacy (e.g., binding affinity). Researchers then submit the top-scoring compounds for experimental validation. Computer docking has proven effective in academic and industrial settings^1,2^.

Neural networks have emerged as a promising method for affinity prediction^3-16^ but with a notable drawback: their predictions often lack interpretability, meaning one cannot easily determine how they arrive at their conclusions. Poor interpretability complicates subsequent lead optimization. For example, neural network scoring functions do not typically indicate which atoms or functional groups contribute most to the overall binding affinity, insight that medicinal chemists could otherwise leverage to improve binding strength and specificity.

We present a novel, interpretable approach for predicting small-molecule binding affinities using context explanation networks (CENs)^17^. Our CENsible scoring function uses deep neural networks to predict the contributions of pre-calculated terms to a given binding affinity rather than directly predicting the affinity itself. In a sense, CENsible is similar to many traditional scoring functions, which predict binding affinities by summing the products of (1) calculated terms (e.g., hydrophobic and steric contributions to binding) and (2) fitted weights (coefficients). However, the CEN-derived weights are not the same for every protein/ligand complex. Instead, CENsible predicts the appropriate weights to apply to the calculated terms based on the specific structure of the protein/ligand complex. CENsible thus leverages the power of machine learning to predict affinity while retaining interpretability; one can easily identify the contribution of each pre-calculated term to the final prediction.

Users can download the CENsible source code free of charge without registration from https://durrantlab.com/censible/, under the terms of the GNU GPL license. The same site links to a helpful Google Colab.

## Materials and Methods

### Data Used for Training and Testing

We downloaded data for training and testing from the PDBbind 2020 database^18-22^, which includes 19,443 crystal structures of protein/ligand complexes taken from the Protein Data Bank with associated experimentally measured binding affinities. We applied several criteria to process this dataset. First, we retained all entries with precisely and approximately defined binding affinities (denoted with “=” and “∼”, respectively). Second, we removed entries with binding affinities measured only as being weaker than a given value (denoted with “>“), judging these to be ambiguously defined. Third, we similarly discarded entries measured only as being stronger than a given value (denoted with “<“) if the associated value was greater than 1 μM; otherwise, these entries were retained. After applying these filters, 19,193 entries remained.

We used a clustered threefold cross-validation scheme to split this data into training and testing sets. We downloaded the Protein Data Bank’s weekly clustering of protein sequences (30% sequence identity)^23-25^ on October 3, 2023. To avoid ambiguity, we removed any multi-chain PDBbind structures belonging to two or more clusters, such that 15,895 PDBbind entries remained. We divided these data into three independent portions, ensuring that entries belonging to the same cluster were always assigned to the same portion. Finally, we trained three different models on two portions, withholding the third as an independent testing set in each case.

### Data Preparation for Training

We standardized all PDBbind protein/ligand structures before training. We removed water molecules and used Open Babel^26^ to add protein and ligand hydrogen atoms appropriate for pH 7. We then voxelized each protein/ligand complex using the *libmolgrid*^27^ python package with default parameters (e.g., resolution of 0.5 Å, 48 × 48 × 48 grid points, 28 atom types; see the Supporting Information for an atom-types list). Voxelization converts a continuous 3D space (e.g., the cartesian coordinates of atomic positions) into a discrete 3D grid well suited as input data for machine-learning models such as convolutional neural networks.

For each PDBbind entry, we also pre-calculated 348 physicochemical terms using *smina*^28^, based on the structure of each protein/ligand complex (see Supporting Information). These included terms used in the AutoDock 4^29^ and AutoDock Vina^30^ scoring functions, terms specific to the *smina* scoring function^28^, counts and whole-ligand features (e.g., ligand length, number of ligand heavy atoms), and a set of 325 steric terms that describe interactions between atoms of specific types (e.g., AliphaticCarbonXSNonHydrophobe vs. OxygenXSAcceptor; *smina* keyword *atom_type_gaussian*). Many of these 325 steric terms were frequently zero; we discarded any that were nonzero in less than 1% of the examples used for training and testing. See the Supporting Information for a complete list of the terms retained in the final scoring function.

The pre-calculated terms had different units and ranges, which could introduce bias into our model. To ensure that all terms were of comparable magnitude, we normalized the data by scaling all terms (i.e., those associated with both training and testing sets):

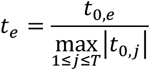

where *t*_*e*_ is the scaled term associated with a given protein/ligand example, *e*; *t*_*0,e*_ is the original unscaled term from *smina*; and *T* is the total number of examples across the training and testing sets. This approach ensured that all terms fell within the range -1 to 1 without changing the sign of any term, which is sometimes physically meaningful.

### Model Training

The CEN scoring functions in the present work use the same deep convolutional neural network architecture as *gnina*’s *default2018* scoring function^14,31,32^, except instead of predicting a single value (binding affinity), they predict a vector of weights (Figure 1). For discussion’s sake, we use *w*_*e*_ to refer to the output weight vector associated with a given protein/ligand example, *e*. These weights serve as coefficients on the scaled pre-calculated terms (*t*_*e*_), such that *w*_*e*_ · *t*_*e*_ approximates the corresponding experimentally measured binding affinity (i.e., pIC_50_, pK_d_, and pK_i_ values derived from the PDBbind 2020 database^18-22^).

**Figure 1.**
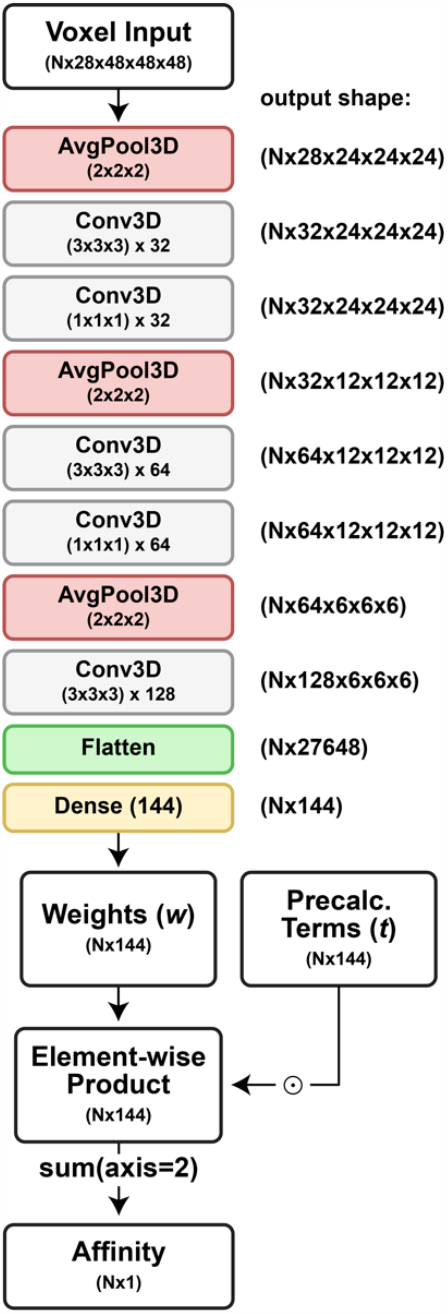
CENsible has the same architecture as *gnina*’s *default2018* model, except the final layer outputs weight vectors (coefficients) rather than affinities. Each Conv3D layer is followed by a rectified linear unit (RELU), not shown. *N* is batch size. To calculate the loss per batch, the model determines per-batch affinity predictions by taking the row-wise dot products of these weights and the precalculated terms.

We trained the CENs for 250 epochs on an NVIDIA GeForce RTX 3090 GPU using the stochastic gradient descent optimizer (learning rate: 0.01; weight decay: 0.0001; momentum: 0.9; see Figure S1) and the StepLR scheduler (step size: 80; gamma: 0.1). Each time a voxel grid was used for training, it was translated by at most 2 Å and randomly rotated. The loss was calculated using the smooth L1 loss criterion^33^, which compared experimental and predicted binding affinities (*w*_*e*_ · *t*_*e*_). The loss was calculated per batch (batch size: 25).

### Model Assessment

We used two methods to assess the predictions of the CEN models. First, we used Pearson’s correlation coefficients to quantitatively assess the accuracy of our model predictions on a withheld test set by comparing predicted and experimentally measured binding affinities.

Second, we used the t-distributed stochastic neighbor embedding (t-SNE) method^34^, implemented in the scikit-learn Python package^35^, to project the CEN-predicted weights onto a two-dimensional space for visualization. K-means clustering verified that similar weight vectors generally map to adjacent regions of this two-dimensional space (Figures S2 and S3). We then identified 916 entries in the PDBbind 2020 database^22^ (used to train CENsible) that also had entries in the SCOP database (release 2022-06-29) with only one unambiguous protein-family assignment. Finally, we identified the SCOP families that were most frequently represented among the labeled PDBbind structures and projected them onto the same t-SNE space with per-family coloring.

### Virtual Screen: *Homo sapiens* pancreatic glucokinase

To confirm CENsible has learned the principles of binding required to separate true ligands from decoys, we performed a virtual screen targeting *Homo sapiens* pancreatic glucokinase, leveraging the files associated with the *HXK4* entry in the DUD-E database^36^. The receptor file associated with this entry was derived from the PDB 3F9M structure^37^. The entry also includes 92 known active molecules, as well as 4,696 decoys (active-to-decoy ratio of roughly 1:51). The DUD-E receptor and compounds already have assigned protonation states and so required no further processing.

We used *smina*^28^ to dock each compound into a box centered on the glucokinase active site. We used *smina*’s “autobox_ligand” parameter to determine the appropriate box dimensions from the DUD-E-provided crystallographic ligand (ligand ID: MRK) and *smina*’s default parameters otherwise. To assess accuracy (e.g., AUROC and EF%), we considered only the top-scoring *smina* pose per ligand, regardless of ionization, tautomerization, etc. We rescored that *smina* pose with our CENsible scoring function.

### Virtual Screen: Influenza Neuraminidase

We also performed a virtual screen targeting influenza neuraminidase, leveraging the files associated with the *NRAM* entry in the DUD-E database^36^. The *NRAM* receptor file was derived from the PDB 1B9V structure^38^. The entry includes 98 known active molecules, as well as 6,200 decoys (active-to-decoy ratio of roughly 1:63). The DUD-E receptor and compounds already have assigned protonation states and so required no further processing.

We used *smina*^28^ to dock each compound into a 20 Å x 20 Å x 20 Å box centered on the neuraminidase active site. We used *smina*’s default parameters otherwise. We again assessed accuracy using AUROC and EF% as above and similarly rescored the *smina* poses with our CENsible scoring function.

### Virtual Screen: *Trypanosoma brucei* methionyl-tRNA synthetase

We also performed a virtual screen targeting *Trypanosoma brucei* methionyl-tRNA synthetase. To prepare the *T. brucei* methionyl-tRNA synthetase structure for docking, we downloaded entry 4EG4^39^ from the Protein Data Bank^24^. We processed the file using the Protein Preparation Workflow available in Schrödinger Maestro 13.5.128 (default parameters). We then removed water and small-molecule-ligand residues and converted S-(dimethylarsenic)cysteine (CAS) residues to cysteine residues.

To prepare small-molecule compounds for the virtual screen, we downloaded the SMILES strings associated with PubChem^40^ bioassay 624268 (primary screen) and bioassay 651971 (confirmatory screen)^41^. We selected the 134 most active compounds from the confirmatory screen (IC_50_ < 5 μM). We also randomly selected 5,360 inactive compounds from the primary screen, such that the active-to-inactive ratio was 1:40. We processed the SMILES strings using Schrödinger’s LigPrep utility to generate 3D structures with enumerated ionization, tautomerization, and chiral states (default parameters, except we generated only at most two stereoisomers per input molecule).

We used *smina*^28^ to dock each compound into a 20 Å x 20 Å x 20 Å box centered on the methionyl-tRNA synthetase active site, using *smina*’s default parameters. We again assessed accuracy using AUROC and EF% as above and similarly rescored the *smina* poses with our CENsible scoring function.

### Tool Use

CENsible is a command-line tool designed to run on Linux-like operating systems. The program itself is written in Python3 (tested on version 3.9.16). It requires the *obabel* (Open Babel) executable to process user-provided protein and small-molecule structures (e.g., to remove water molecules, protonate at pH 7, etc.). It also requires the *smina* executable to calculate *smina* terms for each protein/ligand complex. The git repository includes a helpful README.md file with installation and usage instructions. To encourage broad adoption, we also provide a Google Colab, accessible via https://durrantlab.com/censible/ (see the Supporting Information for a copy of the Python notebook).

### Assistive Writing Technologies

We used assistive writing technologies such as Grammarly and OpenAI’s ChatGPT during manuscript preparation. These supplementary tools acted as editors, not as drivers of content creation. The listed authors thoroughly reviewed, revised, and selectively implemented the suggested edits to ensure accuracy, consistency, and clarity. The responsibility for the paper’s content and quality remains with the authors alone.

## Results and Discussion

In this study, we developed a CEN model for predicting small-molecule binding affinities. Beyond predicting affinities, our approach also indicates the importance of different contributions to those affinities, thus providing an interpretable output that can guide subsequent lead optimization. For each example (*e*) of a protein/ligand complex cataloged in the PDBbind 2020 database^18-22^, we used *smina* to pre-calculate a vector of scaled physicochemical terms, *t*_*e*_. Our model then learned from voxelized representations of the same complexes how to predict a vector of weights, *w*_*e*_, such that the linear sum of these two vectors, *w*_*e*_· *t*_*e*_, approximates the corresponding experimentally measured binding affinity.

### Scoring Function Accuracy

Our primary goal was to develop an *interpretable* machine-learning scoring function, but to have confidence in any such interpretation, the scoring function must also effectively predict ligand binding. To assess the accuracy and consistency of the CEN model, we performed clustered threefold cross validation using 15,895 examples of protein/ligand complexes present in the PDBbind 2020 database. We found that the CEN model effectively predicts binding affinities, with an average testing-set Pearson’s correlation coefficient of 0.5360 (Table 1 and Figure S1). The accuracy of the model was fairly consistent regardless of the training/testing split used (i.e., the standard deviation was small), suggesting (1) accuracy is more a product of learned affinity prediction than the specific data split and (2) the CEN models generalize to new protein classes. A similar analysis showed the CEN approach performs well relative to other scoring functions at assessing crystallographic poses (see Table S1 and related text in the Supporting Information).

**Table 1.** Pearson’s coefficients across three test splits, calculated via a linear fit between known and predicted affinities.

| Split 1 Test | Split 2 Test | Split 3 Test | Mean ± STD |
| --- | --- | --- | --- |
| 0.5661 | 0.5543 | 0.4876 | 0.5360 ± 0.0423 |

Having shown that the three CENs have comparable accuracy regardless of the training/testing split chosen, we trained the final production model on all the PDBbind 2020 data (19,443 entries), which we call the CENsible scoring function. The output vector of this scoring function includes 144 weights to be applied to 144 pre-calculated terms (see Supporting Information). All subsequent analyses leveraged this production model.

### CENsible Tailors Scoring-Function Weights to Specific Targets

We further hypothesized that if CENsible had truly learned to tailor scoring-function weights to binding-pocket properties, similar proteins should have similar predicted weight vectors (*w*_*e*_). To test this hypothesis, we identified SCOP families^42^ that were well represented among PDBbind 2020^18-22^ structures. Proteins belonging to the same SCOP family likely have similar binding pockets because, by definition, they must have at least 30% sequence identity, or some sequence identity in the context of very similar function^42^.

To assess whether proteins of the same family have similar CENsible-predicted *w*_*e*_ vectors, we projected the *w*_*e*_ vectors of 916 SCOP-classified PDBbind structures onto a two-dimensional space using the t-distributed stochastic neighbor embedding (t-SNE) method^34^. We separately colored the structures belonging to well-represented SCOP families to highlight their distribution in this space.

Most of the SCOP families examined had predicted weights that were generally adjacent to each other in single “islands” in t-SNE space, including (1) protein kinases catalytic domain-like, (2) retroviral protease (retropepsin), (3) phosphate binding protein-like, (4) SH3-domain, (5) purine and uridine phosphorylases, and (6) fatty acid binding protein-like (Figure 2). This suggests that CENsible was fairly consistent in its weight predictions when assessing protein/ligand complexes from these families.

**Figure 2.**
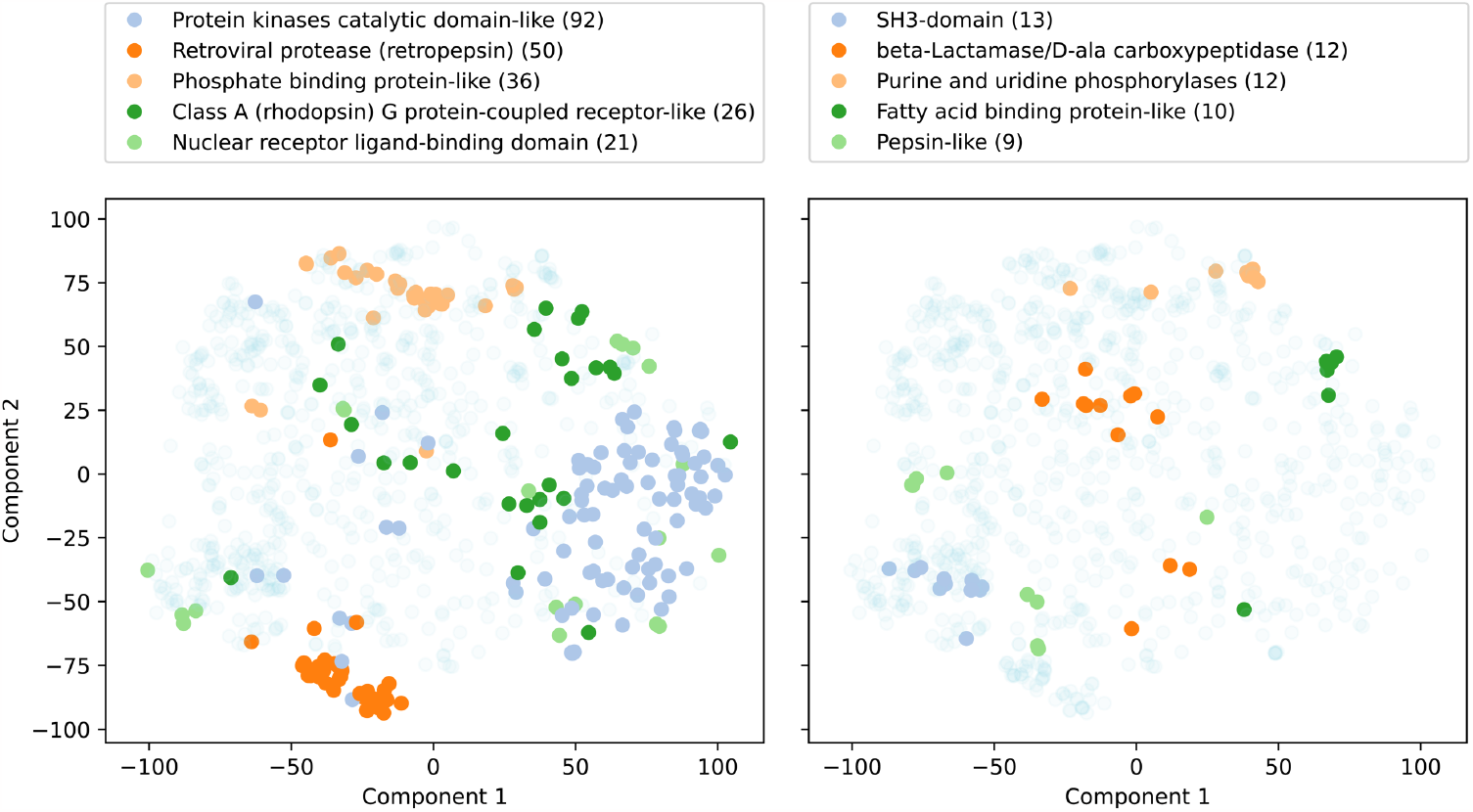
CENsible-predicted term weights (*w*_*e*_) projected onto a 2D t-SNE space, colored by SCOP family label. The most-represented SCOP labels are colored with opaque markers; the remaining labels are colored with transparent markers. The top-five most represented SCOP families are shown in the left panel, and the next five are shown in the right panel. Family names are given above each panel, with the number of examples in parentheses.

The predicted weights of four SCOP families were arguably more dispersed: (1) class A (rhodopsin) G protein-coupled receptor-like, (2) nuclear receptor ligand-binding domain, (3) beta-lactamase/D-ala carboxypeptidase, and (4) pepsin-like (Figure 2). These examples tended to congregate in multiple islands within the t-SNE space rather than one. While it is possible that CENsible has simply not learned to predict consistent *w*_*e*_ vectors for these families, we note that it is also possible for similar vectors to be distant when projected onto t-SNE space in some cases.

These results suggest that CENsible generally produces customized and unique sets of weights for each protein/ligand complex based on the unique features learned from the voxelized representations of the complex. Similar protein/ligand complexes tend to have similar (albeit not identical) weights, and different protein/ligand complexes tend to have different weights.

### Virtual Screening Performance

To assess whether CENsible’s learned principles of ligand binding apply to docked poses, we applied the new scoring function to three virtual screens targeting *H. sapiens* pancreatic glucokinase, influenza neuraminidase, and *T. brucei* methionyl-tRNA synthetase, respectively. In the cases of *H. sapiens* pancreatic glucokinase and influenza neuraminidase, we used known actives and decoy compounds (presumed inactives) cataloged in the DUD-E database^36^. In the case of *T. brucei* methionyl-tRNA synthetase, we used a compound library of known active and inactive compounds identified via PubChem^40^.

We used *smina*^28^ to dock the active and inactive/decoy molecules associated with each protein into the respective active sites and rescored the *smina*-docked poses using our CENsible scoring function. We selected these screens because CENsible effectively prioritized known ligands over other compounds, suggesting it had adequately learned binding principles for these proteins (Figure 3).

**Figure 3.**
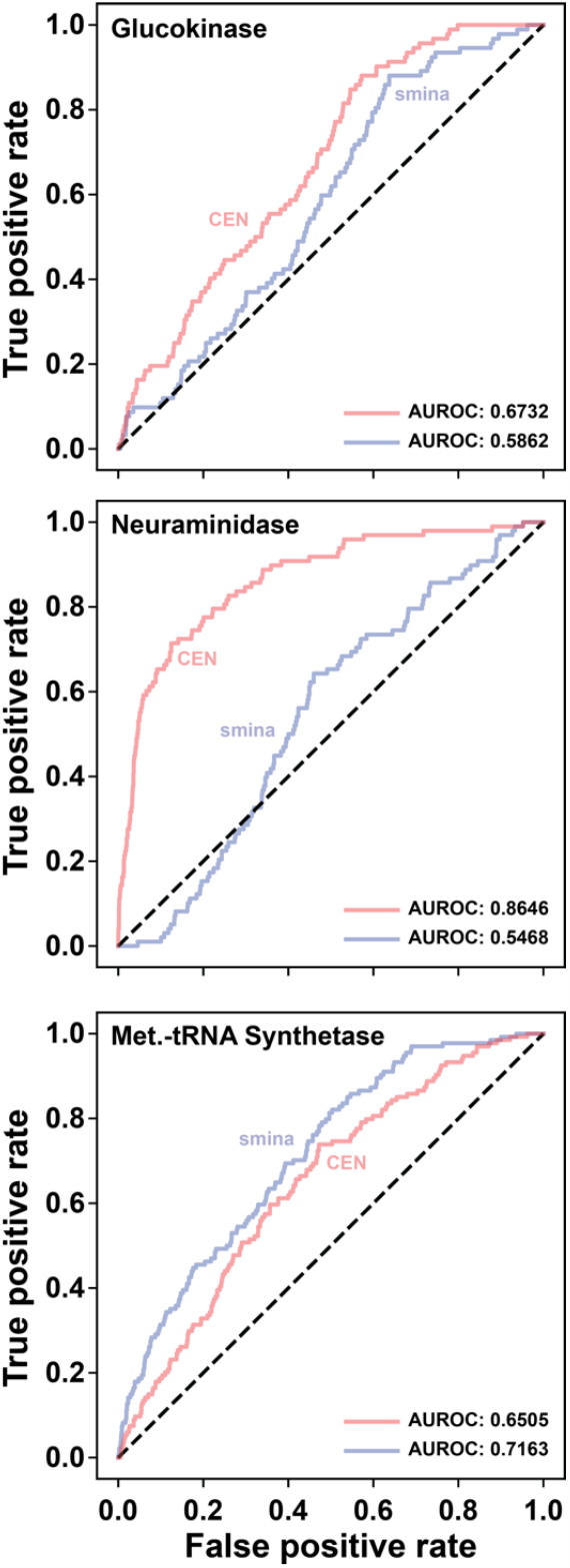
ROC curves associated with the *smina* (blue) and CENsible-rescored (red) virtual screens. The area under each curve is labeled as AUROC. The line of no-discrimination (dotted black line), corresponding to a purely random classifier, is shown for reference.

To evaluate the performance of each scoring function, we calculated the area under the receiver operating characteristic (ROC) curves (AUROC). We selected AUROC because it describes predictivity from the best-ranked compound to the worst and is insensitive to imbalanced data. We reason that if CENsible has learned general principles of ligand binding, it should be able to assess binding across a wide range of affinities.

Although our primary goal is interpretability, it is encouraging that, at least for some systems, CENsible can effectively rank docked poses, even though it was trained on crystallographic poses. The performance of the neuraminidase screens was particularly noteworthy (AUROC 0.8846).

Several important caveats are worth mentioning, however. First, CENsible is only effective if the underlying *smina*-docked poses are reasonably accurate (see Figure S4). It is less likely to accurately predict affinities if given incorrect poses, though poor *smina* performance does not necessarily preclude accurate posing (e.g., *smina*’s AUROC for the neuraminidase screen was only 0.5468).

Second, even if the underlying poses are accurate, CENsible is not suited to every system (Figure S5). As others have noted, the same is true of any docking scoring function^15,18,43-45^. For those systems where CENsible cannot effectively predict affinity, we do not recommend further using it to gain interpretable insights into binding.

Third, even when CENsible is effective at affinity prediction, it is not necessarily the most accurate scoring function for a given system. For example, rescoring the *smina* poses from the glucokinase and methionyl-tRNA synthetase screens with the *gnina default* scoring function gave ROCAUC values of 0.8120 and 0.7225, higher than those obtained when rescoring with CENsible (0.6732 and 0.6505). In contrast, CENsible performed somewhat better than *gnina default* on the neuraminidase screen (0.8646 vs. 0.8126). Further, if the goal is to identify candidate ligands for subsequent testing, top-compound enrichment factors (i.e., the extent to which the top-ranking compounds are enriched with true binders) are arguably more useful metrics for virtual-screen predictivity. By this metric, even *smina* outperforms CENsible in some cases, though in others, CENsible is the clear winner (e.g., the glucokinase and neuraminidase screens; Figure S6). Regardless, CENsible’s primary advantage lies in its ability to provide *interpretable* output tailored to a specific protein/ligand complex.

### Neuraminidase: An Example of Interpretability

Because CENsible predicts weights to apply to pre-calculated terms rather than affinity directly, it offers valuable insights into molecular recognition that can guide subsequent lead optimization. For example, suppose a binding pocket contains a notable hydrophobic sub-pocket. Ligands with hydrophobic moieties positioned in that sub-pocket should have better scores than other compounds, and a scoring function that emphasizes hydrophobic contacts may better prioritize such molecules. In such a case, the CENsible-predicted weights on hydrophobic terms should ideally be higher, indicating that medicinal chemists should further consider hydrophobicity during lead optimization.

To provide a concrete illustration, we compared the weights predicted for the neuraminidase-docked compounds to those predicted for all PDBbind complexes in our training/testing sets. For reference, we first calculated the average predicted weights across the whole PDBbind dataset, with standard deviations (see Supporting Information). We then calculated the average weights across all neuraminidase-docked compounds. Finally, we calculated a z-score for each averaged neuraminidase weight with regard to the PDBbind reference.

Several neuraminidase weights with sizable z-scores suggest receptor-specific compound-optimization strategies. For example, many neuraminidase inhibitors (e.g., oseltamivir; Figure 4) have carboxylate groups that form electrostatic and hydrogen-bond interactions with ARG292, ARG371, and ARG118 (2HU4 numbering^46^). Indeed, CENsible tended to weigh electrostatics more heavily (Table 2). Many neuraminidase inhibitors also have moieties that bind in a hydrophobic subpocket lined in part by ILE222 (e.g., oseltamivir’s pentan-3-yloxy moiety; Figure 4). CENsible also tended to weigh the number of hydrophobic atoms more heavily when predicting neuraminidase inhibitors, perhaps to benefit compounds that could take advantage of this subpocket. This analysis suggests a ligand-optimization strategy that enhances (or at least preserves) these critical electrostatic and hydrophobic interactions.

**Table 2.**
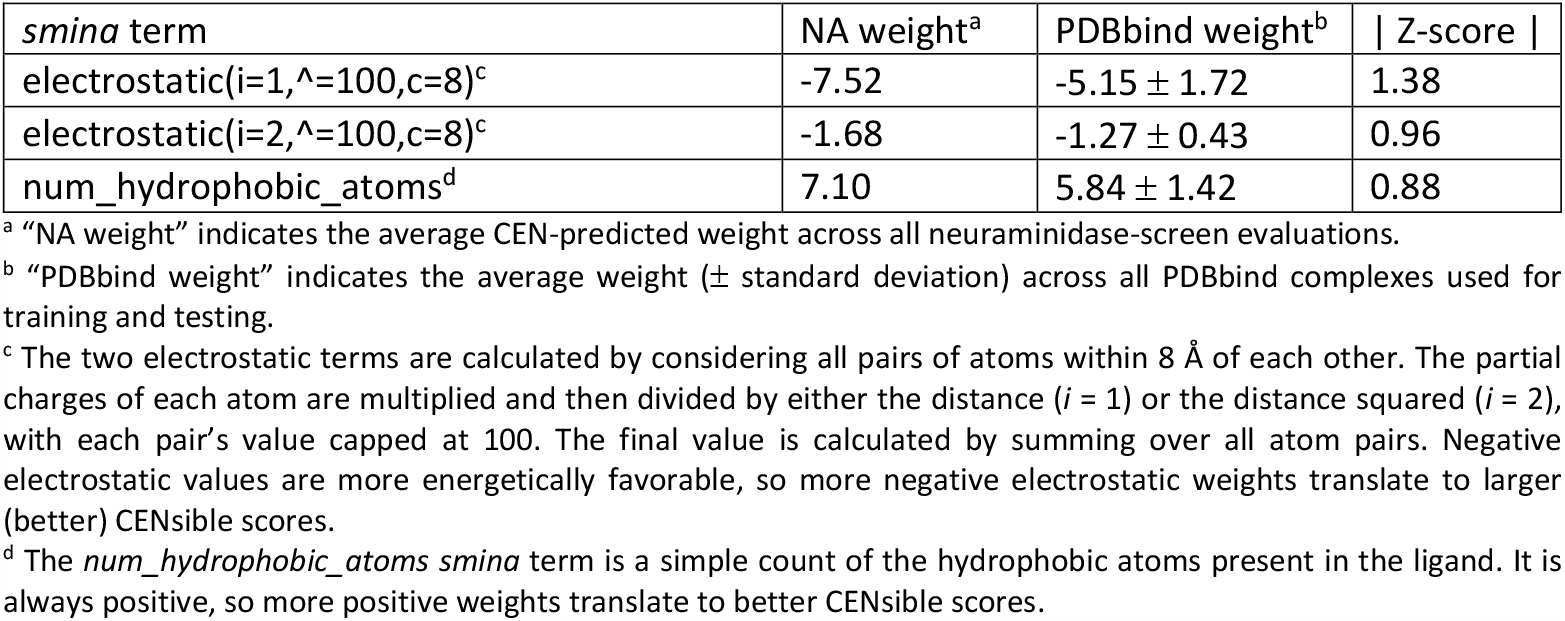
Select *smina* terms whose associated averaged predicted neuraminidase weights differ substantially from those of the entire PDBbind set.

**Figure 4.**
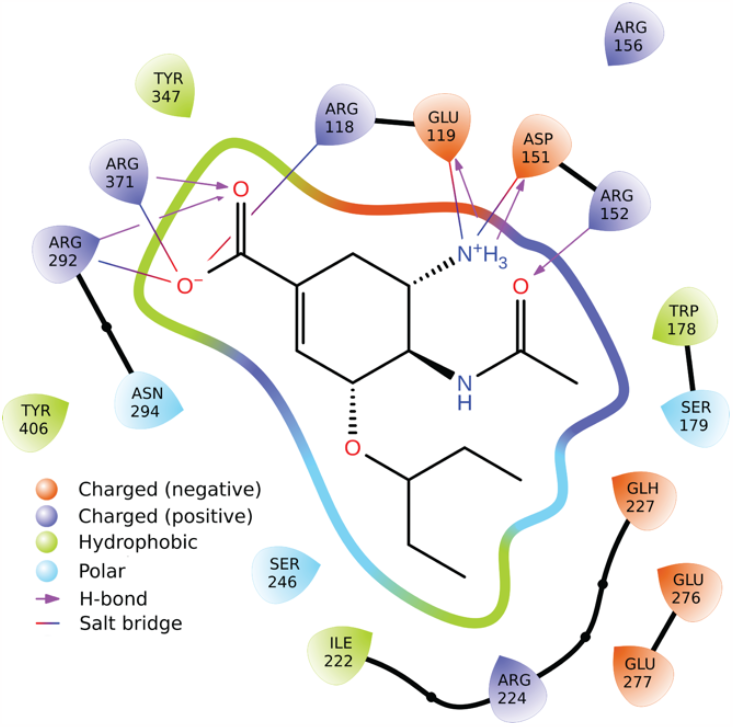
Interactions between influenza neuraminidase and the small-molecule ligand oseltamivir (PDB 2HU4), calculated using Schrödinger’s Ligand Interaction Diagram tool in Maestro.

### Visualizing the Per-Atom Contributions of the Gaussian Steric Terms

Of the 144 pre-calculated *smina* terms that CENsible considers, 123 are pairwise steric (*atom_type_gaussian*) terms. In brief, for each pair of receptor/ligand atoms within 8 Å of each other, *smina* calculates the (1) interatomic distance, *d*; and (2) the “optimal distance,” *d*_*0*_ (i.e., the summed radii of the two atoms). It then assigns a value, *g*, to that pair according to the formula:

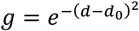

For each combination of receptor/ligand atom types, *smina* calculates the final *atom_type_gaussian* term by summing the associated atom-pair *g* values.

Because these terms are among those directly attributable to specific pairs of receptor/ligand atoms, it is possible to visualize the impact of each atom on each of CENsible’s *atom_type_gaussian*-associated contributions. We calculate the *g* value for each pair, scale that value by the same factor CENsible applies to the corresponding pre-calculated *smina* term, and multiply by the associated CENsible-predicted weight. Finally, we assign half of this *g* value to the receptor atom and half to the ligand atom. Where a single atom contributes to multiple atom-type pairs, the associated values are summed.

We applied this analysis to the same neuraminidase example above. The most beneficial contribution was associated with the *NitrogenXSDonor-OxygenXSAcceptor atom_type_gaussian* term, which contributed +1.05 to CENsible’s final score of 7.45. As shown in Figure 5A, this term captures the same electrostatic interactions seen above (i.e., interactions between the oseltamivir carboxylate group and ARG292, ARG371, and ARG118, as well as interactions between the oseltamivir amine and ASP151 and GLU119), further boosting the influence of electrostatics on the final score (see Table 2).

**Figure 5.**
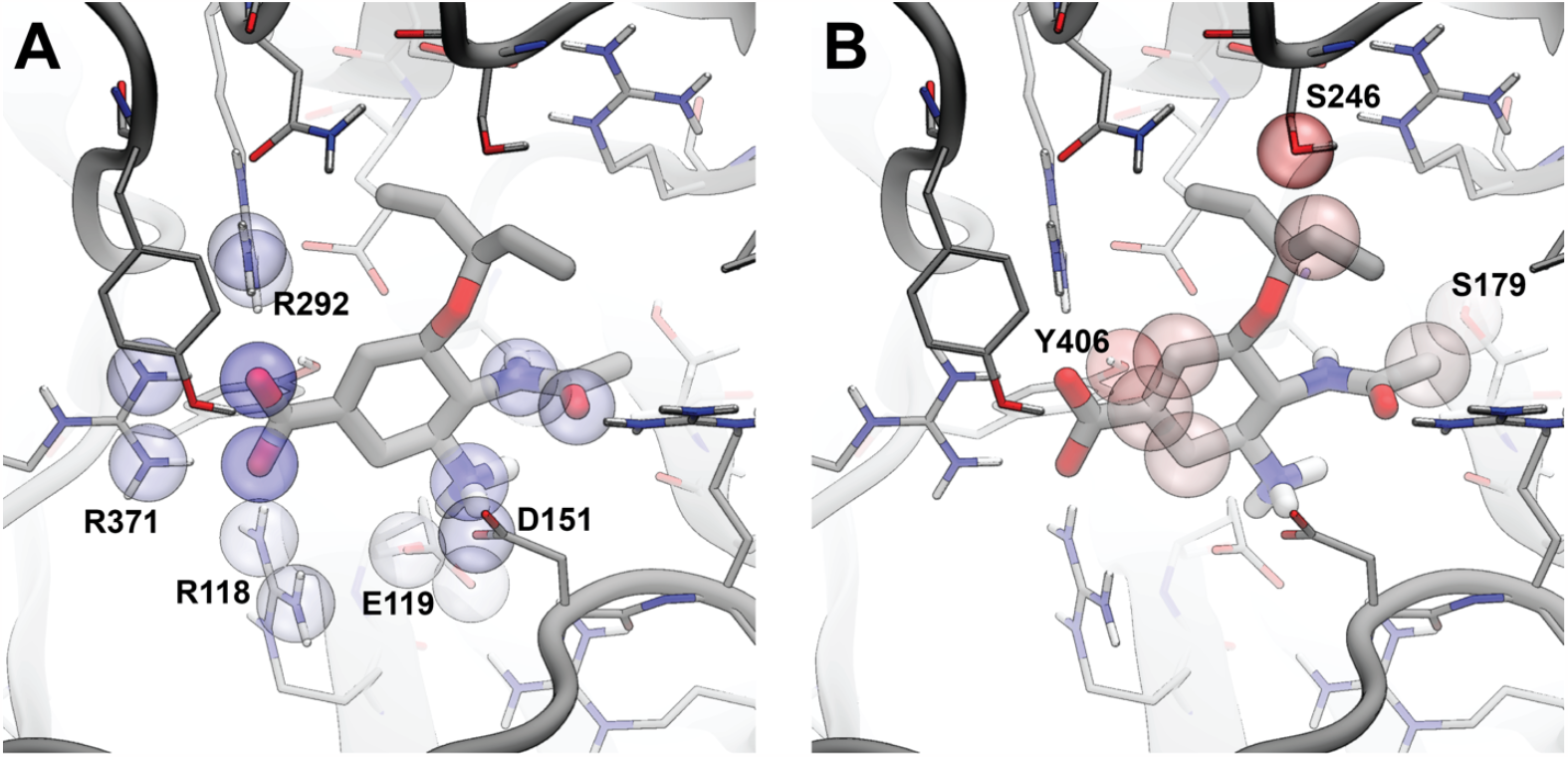
An illustration of the interactions between neuraminidase and oseltamivir. Panel A shows the contributions of beneficial protein/ligand contacts that participate in the *NitrogenXSDonor-OxygenXSAcceptor atom_type_gaussian* term. Panel B shows the contributions of detrimental contacts associated with the *AliphaticCarbonXSHydrophobe*-*OxygenXSDonorAcceptor* term. To simplify the presentation, we highlight only receptor atoms that come within 4 Å of the ligand. We further highlight only those receptor/ligand atoms that contribute more than +0.025 or less than -0.025 to the final CENsible score. Colors range from red to white to blue over the numerical range -0.15 to 0.0 to +0.15. Values outside that range were set to the nearest range boundary.

The *AliphaticCarbonXSHydrophobe-OxygenXSDonorAcceptor atom_type_gaussian* term had the most detrimental impact, decreasing CENsible’s final score by -0.507. Biochemically, it makes sense that this term would have a negative effect, as the juxtaposition of hydrophobic carbon atoms and hydrophilic hydrogen-bond donor/acceptor groups is typically unfavorable. Notably, the SER246 hydroxyl group is near oseltamivir’s pentan-3-yloxy moiety, and the TYR406 hydroxyl group is near its central carbon ring (Figure 5B). Additionally, SER179 is near an oseltamivir methyl group. These findings suggest that modifying oseltamivir to better form hydrogen bonds with these residues could enhance affinity.

Though this analysis technique is powerful, we recommend some caution because CENsible relies in part on overlapping *smina* terms. For example, when scoring potential neuraminidase ligands, CENsible predicted a relatively large weight for the *num_hydrophobic_atoms* term (Table 2), a term that depends only on ligand atoms. However, several *atom_type_gaussian smina* terms also capture hydrophobicity (e.g., the *AliphaticCarbonXSHydrophobe*-*AliphaticCarbonXSHydrophobe* term). Other types of interactions are similarly associated with overlapping, though not identical, terms. Using the *atom_type_gaussian* terms alone to assess the importance of such interactions is thus ill-advised, given that much of their contribution could reside in other terms. Indeed, the summed contribution of all *atom_type_gaussian* terms to a final CENsible score is often only a fraction of the total (e.g., 1.06/7.45 for the neuraminidase/oseltamivir complex).

## Conclusion

We present a novel approach for predicting protein/ligand binding affinities using a context explanation network (CEN). To our knowledge, this study describes the first time CENs have been applied to protein/ligand scoring. Like other machine-learning approaches, our method has effectively learned the principles of ligand binding. However, our model is far more interpretable and so can help guide subsequent lead optimization. We release our software under the GNU GPL license, available for download from https://durrantlab.com/censible/ free of charge, without registration.

## Supporting information

GridAtomTypes.pdf

PDBbind_predicted_weights.xlsx

Precalculated_terms.xlsx

CENsible.ipynb

## Supporting Information

- CENsible_SI.docx: Additional information regarding training, weight-vector clustering, and the CENsible-rescored virtual screens, including Figures S1 – S6.
- CENsible.ipynb: A Google Colab Python notebook for rescoring protein/ligand complexes using CENsible.
- GridAtomTypes.pdf: The receptor and ligand atom types used to generate the voxel grids.
- PDBbind_predicted_weights.xlsx: The CENsible-predicted weights of all PDBbind protein/ligand complexes used for training and testing.
- Precalculated_terms.xlsx: All *smina* terms considered, as well as those included in the final CENsible scoring function.

## Acknowledgments

This work was supported by the National Institute of Health (R01GM132353 and R35GM140753) and the University of Pittsburgh’s Center for Research Computing, RRID:SCR_022735 (supported by NSF OAC-2117681). The content is solely the responsibility of the authors and does not necessarily represent the official views of the National Institutes of Health or the National Science Foundation. The funders had no role in the study design, data collection and analysis, decision to publish, or preparation of the manuscript.

