## Supplementary Information for "CENsible: Interpretable Insights into Small-Molecule Binding with Context Explanation Networks"

### Training Evaluation

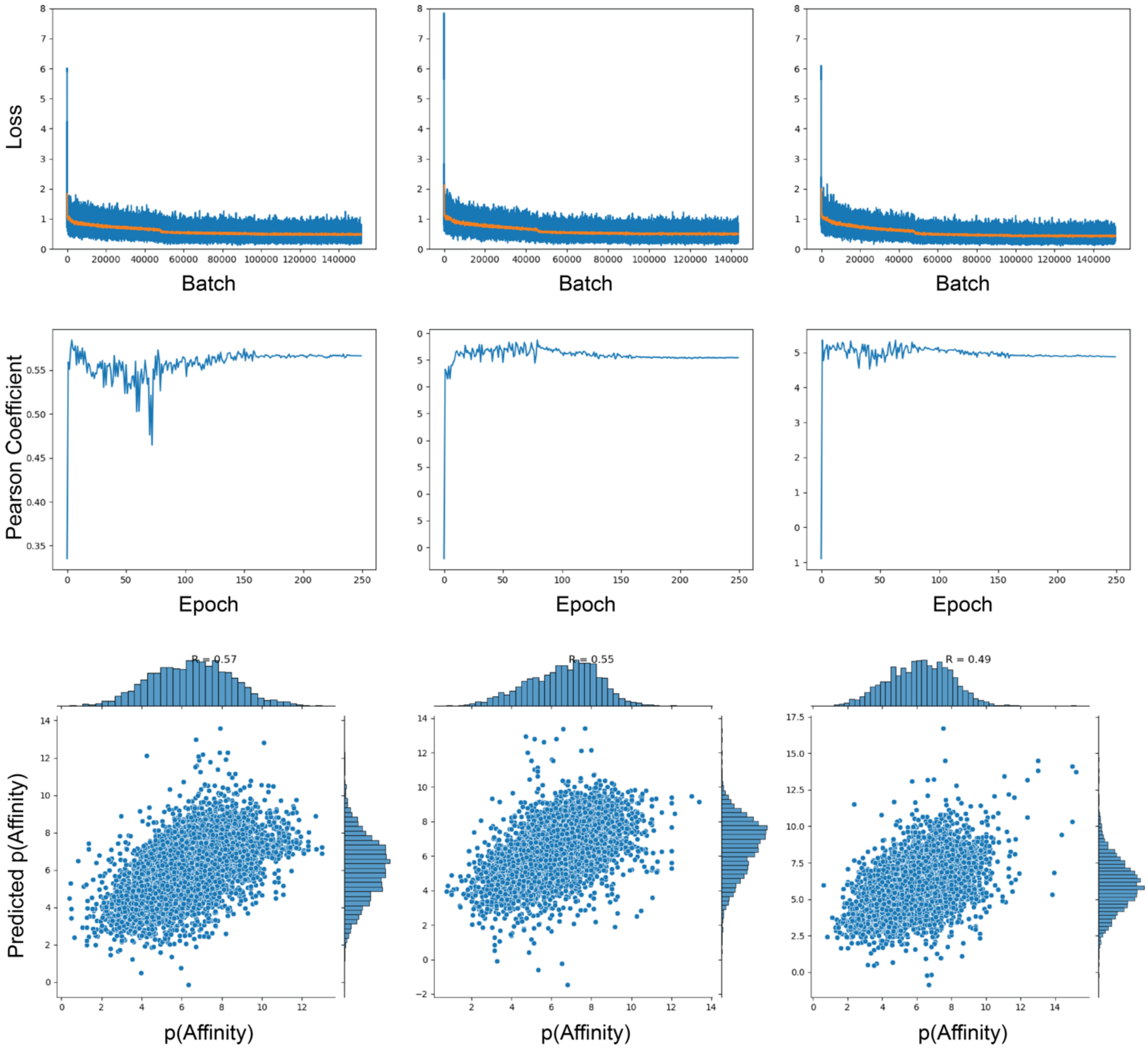


Figure S1. Training evaluations for the first (left), second (middle), and third (right) folds when using clustered cross-validation. Top row: the loss per batch. The orange line represents a moving average of window size 100. Middle row: the Pearson coefficient on the withheld test set for each fold, per epoch of training. Bottom row: a scatter plot of experimental vs. CENsible-predicted affinities (negative logarithm transforms).

### Comparison with Other Scoring Functions at Evaluating Crystallographic Poses

Our primary goal was to develop an *interpretable* machine-learning scoring function, but we were also curious how well our approach could score crystallographic poses compared to other related functions (e.g., *smina*[1] and the *dense*, *gnina2018*, *gnina*, *crossdock_default2018*, *redock_default2018*, and *default2017* neural-network scoring functions available via the *gnina* software package[2,3]).

We could not use the *clustered* cross-validation approach described in the main text to create independent training and testing sets for these other scoring functions because we do not know which proteins were included in their training sets. To ensure a fair comparison, we therefore trained three new CEN models using *unclustered* threefold cross validation. Specifically, we randomly split the PDBbind data into three folds without regard for sequence similarity. We trained three new CEN models on different combinations of two portions, withholding the third as a testing set in each case.

Table S1 shows that the three CEN scoring functions performed better than *smina*. This comparison is critical because the CEN scoring functions incorporate some *smina* terms. The difference is that *smina* uses a single set of weights for all protein/ligand complexes, but the CENs essentially customize the weights based on the structure of the protein/ligand complex.

We also evaluated the same poses present in each cross-validation testing set using *gnina* neural-network-based scoring functions[2,3]. Table S1 shows that the CENs are substantially better at predicting the binding affinities of crystallographic protein/ligand complexes than even the best of the *gnina* scoring functions. Given these results, it is tempting to conclude that the CEN scoring functions are better suited for prospective virtual screens than *gnina* functions. We note, however, that the best performing *gnina* function (*gnina default*) is an ensemble of five models, some of which were trained on docked rather than crystallographic data. The *gnina default* function may therefore be more predictive in the context of a virtual screen[2,3]. Regardless, the strength of the CEN approach is that it is far more interpretable.

Table S1. Pearson's coefficients across three test splits, calculated via a linear fit between known and predicted affinities.

| | Split 1 Test | Split 2 Test | Split 3 Test | Mean ± STD |
|---|---|---|---|---|
| *CEN* | 0.7008 | 0.6868 | 0.6876 | 0.6917 ± 0.0079 |
| \|*smina*\| | 0.2516 | 0.2995 | 0.3275 | 0.2929 ± 0.0384 |
| *gnina default* | 0.5674 | 0.5472 | 0.5555 | 0.5567 ± 0.0102 |
| *dense* | 0.5411 | 0.5221 | 0.5321 | 0.5318 ± 0.0096 |
| *gnina2018* | 0.4973 | 0.4873 | 0.4932 | 0.4926 ± 0.0050 |
| *crossdock_default2018* | 0.4816 | 0.4542 | 0.4617 | 0.4659 ± 0.0141 |
| *redock_default2018* | 0.4978 | 0.4741 | 0.4815 | 0.4845 ± 0.0121 |
| *default2017* | 0.4176 | 0.4258 | 0.4169 | 0.4201 ± 0.0049 |

The training data were randomly split into thirds; for each fold, we trained a CEN model on two thirds of the data and tested on the remaining third (top row). For comparison, we also rescored the test-set examples with *smina* and *gnina*-based scoring functions (subsequent rows). In the case of the *smina* scores, we considered the absolute values.

### Similar CENsible Weight Vectors are Generally Adjacent in t-SNE Space

The t-SNE analysis presented in the main text is predicated on the assumption that similar predicted weight vectors, $w_e$, continue to be adjacent when projected into t-SNE space. To verify this assumption, we first used K-means clustering to identify groups of predicted weight vectors that are similar. We separately divided the data into 2, 3, 4, and 5 clusters and calculated silhouette coefficients for each clustering. These coefficients indicate the relative distance between a given example and its nearest neighbor not in the same cluster, such that high silhouette values suggest better cluster separation. We found that the best separation occurred when we clustered the data into two or three clusters (Figure S2).

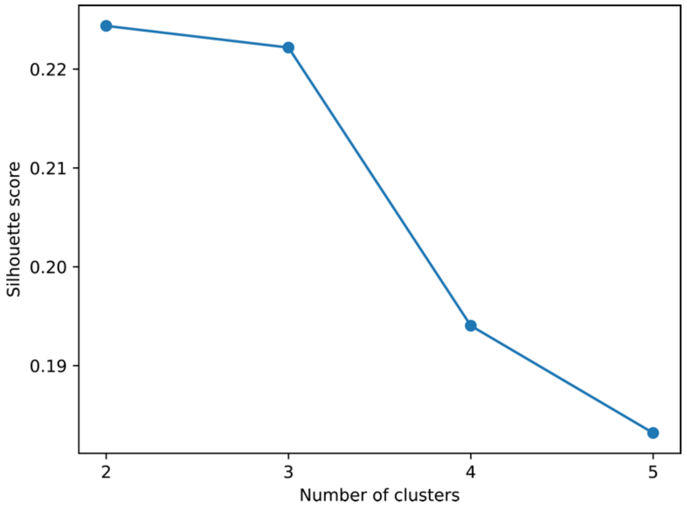


Figure S2. Averaged silhouette coefficient values when generating 2, 3, 4, and 5 clusters.

We next projected the members of these clusters into 2D t-SNE space to determine whether predicted weights belonging to the same cluster continued to be adjacent in the t-SNE space. Adjacency was largely preserved, especially when using two or three clusters (Figure S3, upper row), but also when dividing the data into 4 and 5 clusters (Figure S3, remaining panels). The analysis confirms that clusters are spatially correlated in the lower-dimensional space.

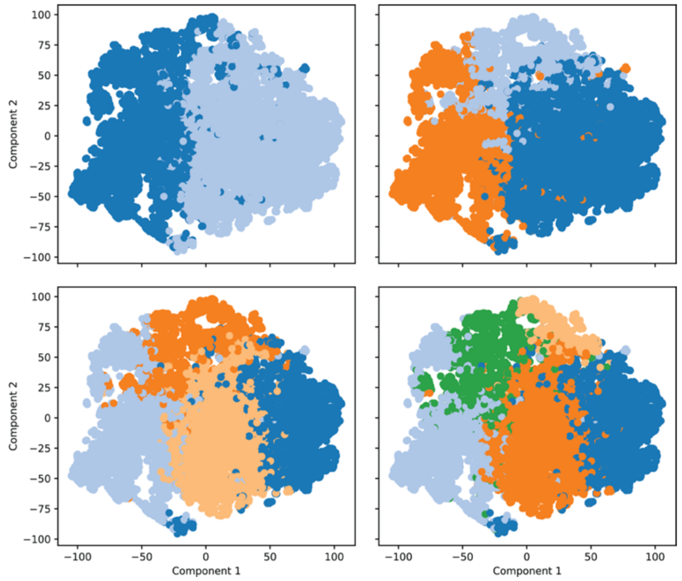


Figure S3. Predicted term weights projected onto 2D t-SNE space, colored by K-means cluster. Colors are randomly assigned.

### CENsible Scores Depend on Accurate Poses

To provide evidence that CENsible scores depend on the accuracy of the underlying ligand poses, we considered all the known-active compounds from the three virtual screens presented in the main text: glucokinase, neuraminidase, and methionyl-tRNA synthetase. We rescored these compounds' best-scoring and worst-scoring *smina* poses with CENsible. We hypothesize that CENsible leverages information about the protein/ligand complex to make its assessments, so the CENsible scores associated with the worst-scoring (likely incorrect) poses should be less than those associated with best-scoring (more likely to be correct) poses.

As shown in Figure S4, this is indeed the case. We used SciPy's[4] *stats.mannwhitneyu* function to perform three two-sided Mann-Whitney *U* tests[5] to assess whether the differences between rescored best/worst *smina* poses are statistically significant. We selected this approach because Shapiro–Wilk tests[6] suggested that some of the score distributions shown in Figure S4 are not normally distributed, complicating the use of a straightforward Student's *t*-test[7]. Per this assessment, the median values are statistically different in all three cases. Further, in all three cases, the median score of the rescored best-scoring *smina* pose was higher (better) than that of the rescored worst-scoring *smina* pose.

These results suggest that CENsible scores improve when reassessing accurate poses, further implying that CENsible's assessment does depend on the structure of the protein-ligand complex.

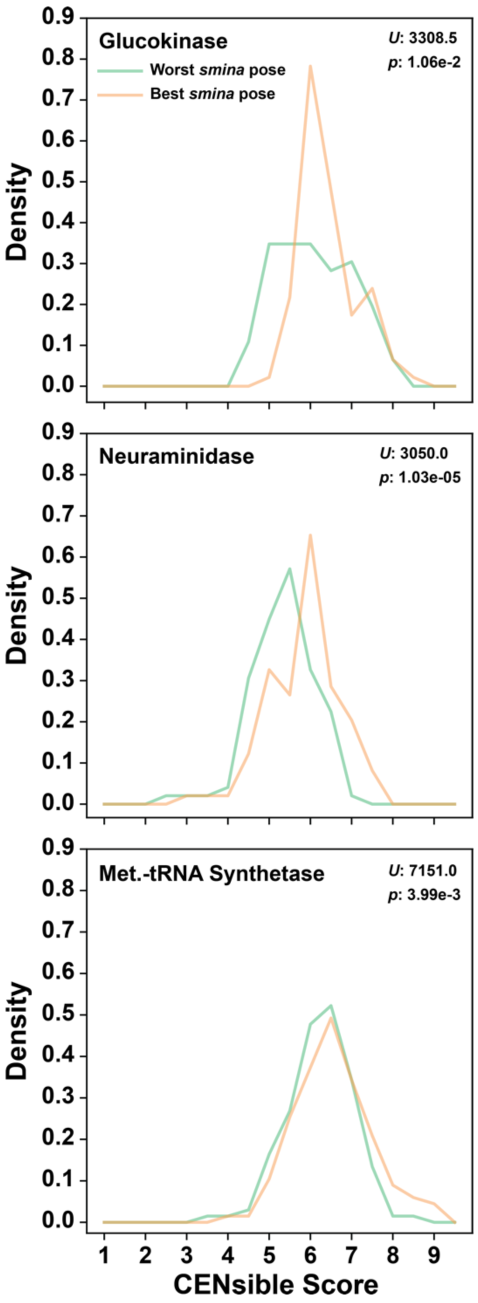


Figure S4. Probability distributions of CENsible-rescored *smina* poses. Only known active compounds from the three virtual screens are included. Bins are spaced 0.5 units, though only every other bin is marked on the X axis. The score distributions associated with the worst and best *smina*-docked poses are shown in green and orange, respectively. *U* statistics and *p* values are given in the upper-right corner of each panel.

### CENsible is not Well Suited to All Systems

Though CENsible generally performs better than *smina* (Pearson's coefficients on testing sets; Table 1), it is not well suited to all systems, as is generally the case with all scoring functions[8-12]. A virtual screen targeting HIV integrase illustrates this variability. The protein and compounds for this screen were again sourced from the DUD-E dataset (id *HIVINT*). The foundational *smina* screen was notably predictive (AUROC: 0.7864; Figure S5). However, when the *smina* poses were reassessed using CENsible, there was a marked decline in the AUROC (0.4949; Figure S5). Similarly, rescoring the poses with the default *gnina* scoring function[2,3] also reduced the AUROC considerably (0.5237, data not shown).

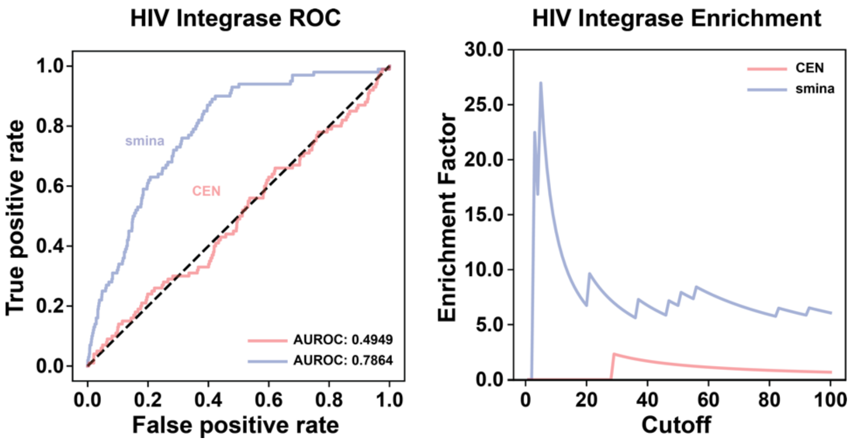


Figure S5. A virtual screen targeting HIV integrase illustrates that CENsible, like all scoring functions, is not well suited to every receptor. On the left, the ROC curves associated with the *smina* (blue) and CENsible-rescored (red) virtual screens. The area under each curve is labeled as AUROC. The line of no-discrimination (dotted black line), corresponding to a purely random classifier, is shown for reference. On the right, the top-compound enrichment factors (i.e., the extent to which the top-ranking compounds are enriched with true binders).

### Virtual Screen Enrichment Factors

In the main text, we use three benchmark virtual screens to assess whether CENsible's learned principles of ligand binding apply to docked poses. We reason that if CENsible has truly learned general principles of ligand binding, it should be able to assess binding across a wide range of affinities. We thus used the AUROC metric to assess these screens (Figure 3) because it captures predictivity from the best-ranked compound to the worst.

Enrichment factors (EFs) are also useful for assessing virtual-screen predictivity. These factors capture the extent to which the top-ranked compounds are enriched with true binders. The EF metric is useful when the goal is not to interpretably assess ligand binding but rather to identify novel ligands. Though EFs are not ideally suited for our assessment, we include them here for completeness' sake.

By the EF metric, CENsible performs well relative to *smina* on the glucokinase and (especially) the neuraminidase screens, but *smina* has better enrichment in the *T. brucei* methionyl-tRNA synthetase screen. Regardless, CENsible's primary utility lies in its ability to provide interpretable output tailored to a specific protein receptor.

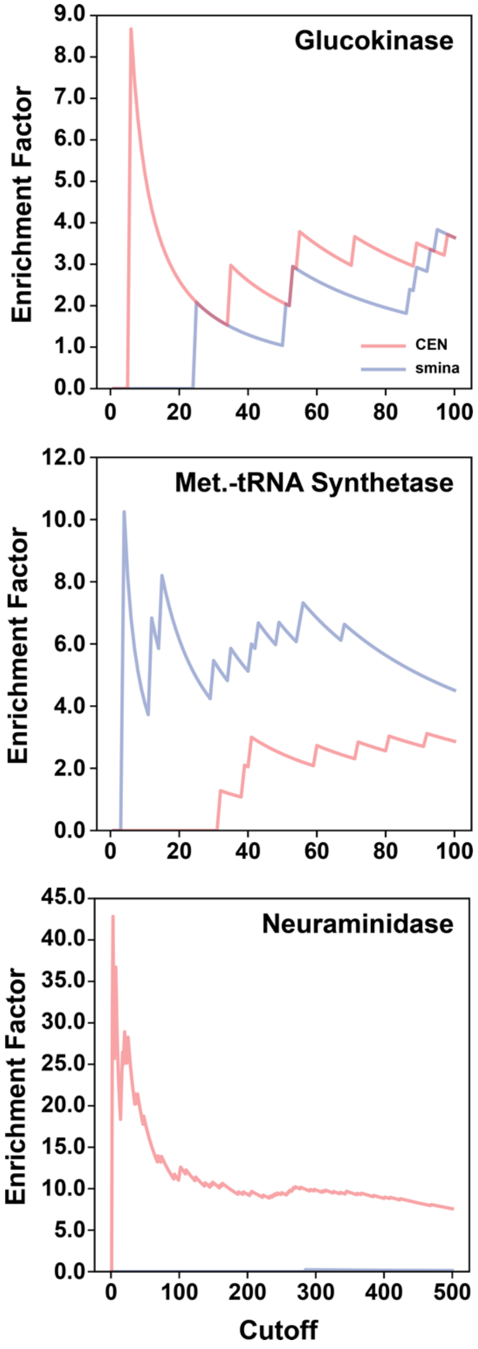


Figure S6. Top-compound enrichment factors (i.e., the extent to which the top-ranking compounds are enriched with true binders).
