## Supplementary material for "CENsible: Interpretable Insights into Small-Molecule Binding with Context Explanation Networks": GridAtomTypes.pdf

### Grid Atom Types

#### Receptor

- AliphaticCarbonXSHydrophobe
- AliphaticCarbonXSNonHydrophobe
- AromaticCarbonXSHydrophobe
- AromaticCarbonXSNonHydrophobe
- Bromine_Iodine_Chlorine_Fluorine
- Nitrogen_NitrogenXSAcceptor
- NitrogenXSDonor_NitrogenXSDonorAcceptor
- Oxygen_OxygenXSAcceptor
- OxygenXSDonorAcceptor_OxygenXSDonor
- Sulfur_SulfurAcceptor
- Phosphorus
- Calcium
- Zinc
- GenericMetal_Boron_Manganese_Magnesium_Iron

#### Ligand

- AliphaticCarbonXSHydrophobe
- AliphaticCarbonXSNonHydrophobe
- AromaticCarbonXSHydrophobe
- AromaticCarbonXSNonHydrophobe
- Bromine_Iodine
- Chlorine
- Fluorine
- Nitrogen_NitrogenXSAcceptor
- NitrogenXSDonor_NitrogenXSDonorAcceptor
- Oxygen_OxygenXSAcceptor
- OxygenXSDonorAcceptor_OxygenXSDonor
- Sulfur_SulfurAcceptor
- Phosphorus
- GenericMetal_Boron_Manganese_Magnesium_Zinc_Calcium_Iron
