## Supplementary material for "CENsible: Interpretable Insights into Small-Molecule Binding with Context Explanation Networks": CENsible.ipynb: CENsible.ipynb

```
# %% [markdown]
# CENsible 1.0

CENsible uses context explanation networks (CENs) to predict small-molecule binding affinities. Rather than predict a binding affinity directly, it predicts the contributions of pre-calculated terms to the overall affinity, thus providing interpretable output. CENsible insights are useful for subsequent lead optimization.

We release CENsible under the terms of the [GNU General Public License
v3.0](https://github.com/durrantlab/censible/blob/main/LICENSE.md). The git repository is available at [https://github.com/durrantlab/censible](https://github.com/durrantlab/censible).

# %% [markdown]
# 1. Install CENsible and Define Colab Helper Functions

# %% [code]
# @title Install CENsible and its dependencies

# 1/ Clone the repo
%cd /content/
!rm -rf /content/censible/
!git clone https://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@github.com/durrantlab/censible.git
%cd /content/censible/

# 2. Install dependencies

# Colab comes with it's own CUDA-enabled pytorch.

# Install the other dependencies (not torch)
!cd /content/censible/ && \
 cat requirements_predict.txt | grep -v torch > requiements_without_torch.txt && \
 cat requiements_without_torch.txt
!cd /content/censible/ && pip install -r requiements_without_torch.txt

# Download smina and open babel
!apt-get install libeigen3-dev
!apt-get install libopenbabel-dev
!cd /content/censible/ && \
    curl -O https://raw.githubusercontent.com/durrantlab/colab-support-files/main/smina
!cd /content/censible/ && chmod a+rwx ./smina

!pip install openbabel-wheel==3.1.1.17
!pip install py3Dmol


# %% [code]
# @title Define functions for use in Colab

import pandas as pd
import py3Dmol

def show_terms_and_weights():
  with open('/content/censible/test_out.tsv', 'r') as f:
      lines = f.readlines()

  data = [line.rstrip().split('\t') for line in lines[14:20]]

  df = pd.DataFrame(data[1:], columns=data[0])
  # print("".join(lines[:14]))

  # return "".join(lines[:14]), df
  return df

def show_pdb_chain_info():
  with open("/content/censible/test_out.pdb") as f:
    remarks = [l[7:] for l in f.readlines()[1:] if l.startswith("REMARK")]
  remarks = [r for r in remarks if "LIGAND" not in r]
  remarks = [r for r in remarks if "RECEPTOR" not in r]
  remarks = [r for r in remarks if "CONTRIBUTION TO SCORE" not in r]
  remarks = [r for r in remarks if "ATOM-TYPE PAIR" not in r]

  chain_defs = {
    # r.split()[0] : ("".join(r.split()[1:]), r.split()[3])
    r.split()[0] : ("".join(r.split()[1:4]), r.split()[4])
    for r in remarks if "      " in r and r.split()[0] != "CHAIN"
  }

  print("")
  print("".join(remarks))

  return chain_defs

def run_censible(lig_path, recep_path, chain_to_show=None):
  # Always regenerate converted receptor file
  !rm -f {recep_path}*converted*

  # Also delete old output files
  !rm -f /content/censible/test_out.tsv /content/censible/test_out.pdb

  !cd /content/censible/ && python predict.py \
      --ligpath {lig_path} \
      --recpath {recep_path} \
      --smina_exec_path /content/censible/smina \
      --obabel_exec_path /usr/local/bin/obabel \
      --tsv_out /content/censible/test_out.tsv \
      --pdb_out /content/censible/test_out.pdb

  # Clean up
  !rm -f /content/censible/*converted*


def read_pdb_by_chain(filename):
    chains = {}

    with open(filename, 'r') as f:
        for line in f:
            # Check if the line is an ATOM or HETATM record
            if line.startswith(("ATOM", "HETATM")):
                chain_id = line[21]
                if chain_id not in chains:
                    chains[chain_id] = []
                chains[chain_id].append(line)

    for chain in chains:
      chains[chain] = "".join(chains[chain])

    return chains

def extract_b_factors(pdb_txt):
    b_factors = []

    # Split the PDB text into lines and iterate over them
    for line in pdb_txt.splitlines():
        # Check if the line is an ATOM or HETATM record
        if line.startswith(("ATOM", "HETATM")):
            # B-factor is found in columns 61-66 (1-based numbering)
            b_value = float(line[60:66].strip())
            b_factors.append(b_value)

    return b_factors

def vis_3d_results(chain_id):
  pdbs_by_chain = read_pdb_by_chain('/content/censible/test_out.pdb')

  view = py3Dmol.view()
  view.setBackgroundColor('white')

  # First add the protein
  view.addModel(pdbs_by_chain["Y"], 'pdb')
  mol = view.getModel()
  mol.setStyle({'chain': 'Y'}, {'cartoon': {'color': 'lightgray'}})
  mol.setStyle({'chain': 'Y'}, {'stick': {"radius": 0.05, 'colorscheme': 'grayCarbon'}}, True)

  # And the chain
  if chain_id != "None (Visualize Only Receptor/Ligand Atoms)":
    view.addModel(pdbs_by_chain[chain_id], 'pdb')
    mol = view.getModel()
    mol.setStyle(
        {'chain': chain_id},
        {'sphere': {
            "radius": 0.75, "opacity": 0.9,
            'colorscheme':{'prop': 'b','gradient': 'RWB','min': -0.5,'max': 0.5}
        }}
    );

  # Now the ligand
  view.addModel(pdbs_by_chain["Z"], 'pdb')
  mol = view.getModel()
  mol.setStyle({'chain': 'Z'}, {"stick": {"radius": 0.4, 'colorscheme': 'grayCarbon'}})

  view.zoomTo({'chain': 'Z'})

  view.show()

# %% [markdown]
# 2. Test CENsible Installation Using Example Files

# %% [code]
# @title Run CENsible on the example receptor/ligand files

run_censible("censible/data/test/1wdn_ligand.mol2", "censible/data/test/1wdn_receptor.pdb")

# %% [code]
# @title Display the terms and contributions leading to the prediction

show_terms_and_weights()

# %% [code]
# @title Get a description of each detected contribution, with an associated PDB chain ID for reference

# Show pdb info
chain_defs = show_pdb_chain_info()


# %% [code]
# @title Visualize the contributions of a given interaction (by Chain ID)
# @markdown Which chain (interaction) from the list above would you like to visualize?
chain = "None (Visualize Only Receptor/Ligand Atoms)" # @param ["None (Visualize Only Receptor/Ligand Atoms)", "A", "B", "C", "D", "E", "F", "G", "H", "I", "J", "K", "L", "M", "N", "O", "P", "Q", "R", "S", "T", "U", "V", "W", "X", "a", "b"]

if chain == "None (Visualize Only Receptor/Ligand Atoms)":
  print("Interaction: None (receptor/ligand atoms only)")
else:
  label, contribution = chain_defs[chain]
  print(f"Interaction: {label}; Contribution: {contribution}")
  print("")

# Visualize one of the chains
vis_3d_results(chain)

# %% [markdown]
# 3. Use CENsible on Your Own Receptor/Ligand Files

# %% [code]
# @title Upload your own files and run CENsible on them
# @markdown Note: You can run this cell as many times as you like to evaluate different protein/receptor affinities.
# @title
from google.colab import files
import os

!rm -f /content/censible/*.tsv /content/censible/receptor.* /content/censible/ligand.*

print("Upload the receptor file:")
recep_file = files.upload()
recep_filename = list(recep_file.keys())[0]
recep_ext = recep_filename.split(".")[-1]
new_recep_filename = "/content/censible/receptor." + recep_ext

with open(new_recep_filename, "w") as f:
  content = list(recep_file.values())[0]
  if isinstance(content, bytes):
      content = content.decode('utf-8')
  f.write(content)

print("\nUpload the ligand file:")
lig_file = files.upload()
lig_filename = list(lig_file.keys())[0]
lig_ext = lig_filename.split(".")[-1]
new_lig_filename = "/content/censible/ligand." + lig_ext

with open(new_lig_filename, "w") as f:
  content = list(lig_file.values())[0]
  if isinstance(content, bytes):
      content = content.decode('utf-8')
  f.write(content)

df = run_censible(new_lig_filename, new_recep_filename)

# Clean up
os.unlink(recep_filename)
os.unlink(lig_filename)
os.unlink(new_recep_filename)
os.unlink(new_lig_filename)

df


# %% [code]
# @title Display the terms and contributions leading to the prediction

show_terms_and_weights()

# %% [code]
# @title Get a description of each detected contribution, with an associated PDB chain ID for reference

# Show pdb info
chain_defs = show_pdb_chain_info()

# %% [code]
# @title Visualize the contributions of a given interaction (by Chain ID)
# @markdown Which chain (interaction) from the list above would you like to visualize?
chain = "None (Visualize Only Receptor/Ligand Atoms)" # @param ["None (Visualize Only Receptor/Ligand Atoms)", "A", "B", "C", "D", "E", "F", "G", "H", "I", "J", "K", "L", "M", "N", "O", "P", "Q", "R", "S", "T", "U", "V", "W", "X", "a", "b"]

if chain == "None (Visualize Only Receptor/Ligand Atoms)":
  print("Interaction: None (receptor/ligand atoms only)")
else:
  label, contribution = chain_defs[chain]
  print(f"Interaction: {label}; Contribution: {contribution}")
  print("")

# Visualize one of the chains
vis_3d_results(chain)
```
